# *In situ* activation and heterologous production of a cryptic lantibiotic from a plant-ant derived *Saccharopolyspora* species

**DOI:** 10.1101/733808

**Authors:** Eleni Vikeli, David A. Widdick, Sibyl F. Batey, Daniel Heine, Neil A. Holmes, Mervyn J. Bibb, Dino J. Martins, Naomi E. Pierce, Matthew I. Hutchings, Barrie Wilkinson

**Author notes:** Address correspondence to Barrie Wilkinson and Matthew I. Hutchings. These authors contributed equally to this work (Eleni Vikeli, David A. Widdick, Sibyl F. Batey).

## Abstract

Most clinical antibiotics are derived from actinomycete natural products (NPs) discovered at least 60 years ago. Repeated rediscovery of known compounds led the pharmaceutical industry to largely discard microbial NPs as a source of new chemical diversity but advances in genome sequencing revealed that these organisms have the potential to make many more NPs than previously thought. Approaches to unlock NP biosynthesis by genetic manipulation of the strain, by the application of chemical genetics, or by microbial co-cultivation have resulted in the identification of new antibacterial compounds. Concomitantly, intensive exploration of coevolved ecological niches, such as insect-microbe defensive symbioses, has revealed these to be a rich source of chemical novelty. Here we report the novel lanthipeptide antibiotic kyamicin generated through the activation of a cryptic biosynthetic gene cluster identified by genome mining *Saccharopolyspora* species found in the obligate domatia-dwelling ant *Tetraponera penzigi* of the ant plant *Vachellia drepanolobium*. Heterologous production and purification of kyamicin allowed its structural characterisation and bioactivity determination. Our activation strategy was also successful for the expression of lantibiotics from other genera, paving the way for a synthetic heterologous expression platform for the discovery of lanthipeptides that are not detected under laboratory conditions or that are new to nature.

**Importance:** The discovery of novel antibiotics to tackle the growing threat of antimicrobial resistance is impeded by difficulties in accessing the full biosynthetic potential of microorganisms. The development of new tools to unlock the biosynthesis of cryptic bacterial natural products will greatly increase the repertoire of natural product scaffolds. Here we report an activation strategy that can be rapidly applied to activate the biosynthesis of cryptic lanthipeptide biosynthetic gene clusters. This allowed the discovery of a new lanthipeptide antibiotic directly from the native host and via heterologous expression.

---

Antimicrobial resistance (AMR) is arguably the greatest health threat facing humanity in the 21^st^ century (1–3). It is predicted that without urgent action, infectious disease will become the biggest killer of humans by 2050 (1). The majority of clinically used antibiotics are based on microbial natural products, isolated mostly from soil-dwelling *Streptomyces* species and other filamentous actinomycete bacteria, and these organisms remain a promising source of new antibiotics. Although the discovery pipeline began to dry up in the 1960s, blighted by the rediscovery of known compounds, we know from large scale genome sequencing that up to 90% of microbial natural products are not produced under laboratory conditions (4). Thus, there exists a wealth of novel chemistry waiting to be discovered by mining the genomes of these organisms. Bearing in mind that >600 *Streptomyces* species and many other so called ‘rare’ actinomycetes have been described, thousands of potentially useful but “cryptic” bioactive compounds are waiting to be discovered, even from well-characterised strains (5,6). Several approaches have been taken to activate cryptic pathways including the heterologous expression of entire biosynthetic gene clusters (BGCs) in optimised *Streptomyces* host strains, and rewiring BGCs to bypass their natural regulatory mechanisms (7). The knowledge that we have barely sampled the biosynthetic capabilities of known strains, and that even well explored environments such as soil have been under sampled for antibiotic-producing microbes, provides a much-needed opportunity for the development of new natural product-based antibiotics.

Searching symbiotic niches for new actinomycete strains also shows great promise for discovering new natural products (8–11). We previously described the formicamycins, new polyketides with potent Gram-positive antibacterial activity produced by a new *Streptomyces* species that we named *Streptomyces formicae* KY5 (12). This species was isolated from a phytoecioius ant species, *Tetraponera penzigi*, whose colonies inhabit the African ant plant *Vachellia (=Acacia) drepanolobium*. The ants were collected in Kenya, hence the KY strain designation (13). These ants live in symbiosis with their host plants, the “whistling thorn acacias”, that have evolved specialised hollow, stipular thorns called domatia to house the ants (14). In return for housing, plant ants protect their hosts against attack by large herbivores, including elephants (15), and recent reports have suggested that they grow specialized fungal communities inside their domatia, possibly as a food source for their larvae (16,17).

The external, cuticular microbiome of *T*. *penzigi* ants is heterogeneous, and unbiased methods have shown this is dominated by members of the phyla Proteobacteria and Firmicutes, with Actinobacteria forming a minor component (13). This contrasts with the better studied fungus-farming leafcutter ants of the tribe Attini, which are dominated by actinobacteria, specifically by a single strain of *Pseudonocardia* that can be vertically transmitted by the new queens (18,19). Leafcutter ants feed cut plant material to their symbiotic food fungus *Leucoagaricus gongylophorus* and use antifungals made by their *Pseudonocardia* symbionts to defend their food fungus against fungal parasites in the genus *Escovopsis* (20–22). Despite the low abundance of actinobacteria, we isolated several strains, including three from the rare actinomycete genus *Saccharopolyspora*, which, despite the modest number of described species, is the origin of the medically and agriculturally important natural products erythromycin and spinosyn.

Genome mining of these *Saccharopolyspora* strains identified a conserved BGC encoding a putative cinnamycin-like lanthipeptide antibiotic (lantibiotic) (23), although no products for this BGC could be identified from the wild-type isolates. Cinnamycin is a class II type B lantibiotic produced by *Streptomyces cinnamoneus* DSM 40005 which destabilises the cytoplasmic membrane by binding phosphatidylethanolamine (PE) (23–25). Lanthipeptides belong to the ribosomally synthesised and post-translationally modified peptide (RiPP) family of natural products (26,27), and cinnamycin is the founding member of a sub-group of lanthipeptide RiPPs with antibacterial activity that includes cinnamycin B (28), duramycin (29), duramycin B and C (30), and mathermycin (31) (Fig. 1A). These molecules are produced by actinomycetes and comprise 19 amino acid residues, several of which are modified to generate lanthionione or methyllanthionine cross-links (26,27). Additional modifications include β-hydroxylation of the invariant aspartic acid residue at position 15 and formation of an unusual lysinoalanine cross-link between the serine residue at position 6 and lysine residue at position 19 (32–34). The interaction of these molecules with PE has therapeutic potential: duramycin binds to human lung epithelial cell membranes leading to changes in the membrane, or its components, promoting chloride ion secretion and clearance of mucus from the lungs (25). On this basis, duramycin entered Phase II clinical trials for the treatment of cystic fibrosis (35).

**FIG 1.**
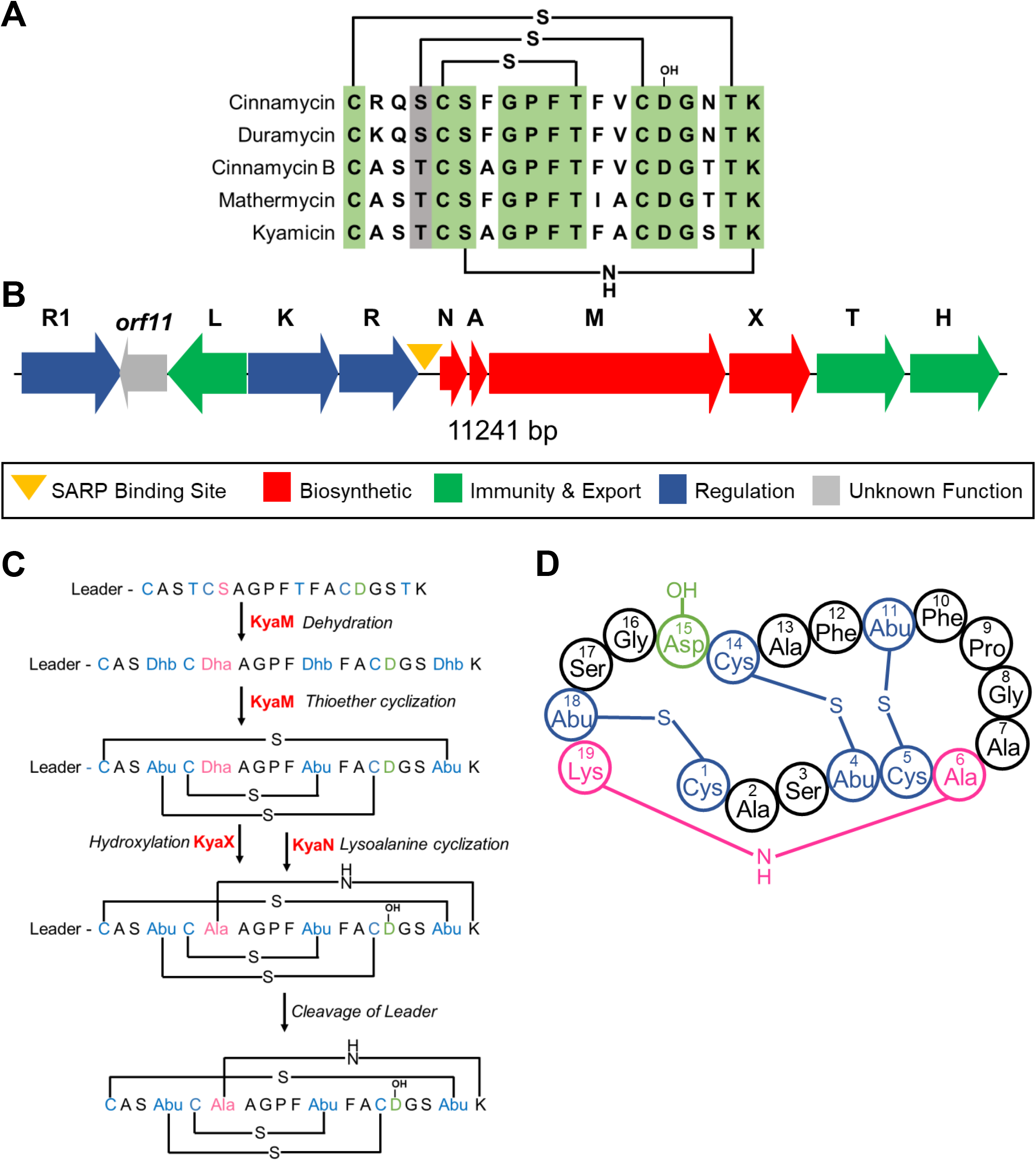
Kyamicin peptide sequence and biosynthesis. **(A)** Alignment of core peptides of kyamicin and a selection of known Type B cinnamycin-like lantibiotics, with the positions of the thioether and lysinoalanine bridges in the mature peptide shown. Conserved residues are highlighted in green, similar residues are highlighted in grey. **(B)** The kyamicin biosynthetic gene cluster, with genes colored according to predicted function. **(C)** Schematic of kyamicin biosynthesis. The thioether bridges are formed first by dehydration of Thr4, Thr11, Thr18 and Ser6 by KyaM to form dehydrobutyrine (Dhb) and dehydroalanine (Dha) residues, respectively. After thioether cyclization by KyaM, Dhb becomes S-linked Abu and Dha becomes S-linked Ala. Asp15 is hydroxylated by KyaX and the lysinoalanine bridge is then formed between Dha6 and Lys19 by KyaN. After the core peptide is fully modified, the leader peptide is proteolytically cleaved. **(D)** Structural representation of the mature kyamicin lantibiotic.

Here we describe activation of the cryptic *Saccharopolyspora* lanthipeptide BGCs and the characterization of their product, a new class II lantibiotic that we called kyamicin. We also exemplify a heterologous expression platform for lanthipeptide production that may be particularly useful for strains that are refractory to genetic manipulation. The methodologies reported should be applicable for the activation of cryptic BGCs from a wide range of actinomycetes.

## RESULTS

### Origin, characteristics and genome sequencing of *Saccharopolyspora* strains

The *Saccharopolyspora* strains were isolated from ants taken from the domatia of *T. penzigi* plant ants collected in two locations in Kenya (13), and named KY3, KY7 and KY21. 16S rDNA was amplified and Sanger sequenced using the universal primers 533F and 1492R (Genbank accession numbers JX306001, JX306003, JX306004, respectively). Alignments show that KY3 and KY7 are identical across the sequenced 16S rDNA region while KY21 differs by a single base pair (Fig. S1). Further analysis showed that all three strains share 99% sequence identity with *Saccharopolyspora* 16S rDNA sequences in public databases. High molecular weight genomic DNA was isolated from each strain, sequenced at the Earlham Institute (Norwich, UK) using SMRT sequencing technology (Pacific Biosciences RSII platform) and assembled using the HGAP2 pipeline as described previously (36). This gave three circular chromosomes of approx. 6.33 Mbp, the full analysis of which will be reported separately. Alignment of the KY3 and KY7 genome sequences using RAST SEED Viewer and BLAST dot plot revealed a full synteny along their genomes with 99-100% sequence identity at the nucleotide level suggesting KY3 and KY7 are the same strain and different to KY21.

### Identification of a conserved cinnamycin-like BGC

The biosynthetic potential of all three strains was probed using the genome mining platform antiSMASH (37). The three genomes each encode approximately 25 BGCs with significant overlap. Amongst these was a BGC for a cinnamycin-like lanthipeptide. The BGC architecture was conserved across all three genomes, including an identical pro-peptide sequence encoded by the precursor peptide gene, suggesting they all encode the same molecule which we named kyamicin (Fig. 1B). The sequence and annotations for these three BGCs have been deposited at GenBank under the accession numbers MK251551 (KY3), MK251552 (KY7) and MK251553 (KY21).

Through comparison to the cinnamycin BGC (24), and cinnamycin biosynthesis (32), we assigned roles to each of the genes in the kyamicin (*kya*) BGC (Table 1). The *kya* BGC is more compact than that for cinnamycin, and the genes missing from the kyamicin BGC are dispensable for cinnamycin production (38). The *cinorf11* gene is not required for cinnamycin production but a homologue is present in the kyamicin cluster. While *cinorf11* lacks a plausible stop codon and its reading frame extends 570 bp into the *cinR1* gene, its homologue, *kyaorf11*, has a stop codon and does not run into the *kyaR1* gene suggesting it may encode a functional protein.

**TABLE 1:** Proteins encoded by the cinnamycin and kyamicin BGCs.

| **Kyamicin** | **Cinnamycin** | **Duramycin** | **Proposed function** |
| --- | --- | --- | --- |
| **KyaN (123aa)** | CinN (119aa) | DurN (119aa) | Formation of lysinoalanine bridge |
| **KyaA (78aa)** | CinA (78aa) | DurA (77aa) | Precursor peptide |
| **KyaM (1065aa)** | CinM (1088aa) | DurM (1083aa) | Formation of lanthionine residues |
| **KyaX (302aa)** | CinX (325aa) | DurX (327aa) | Hydroxylation of Asp15 |
| **KyaT (327aa)** | CinT (309aa) | DurT (352aa) | Export |
| **KyaH (294aa)** | CinH (290aa) | DurH (290aa) | Export |
| **Not Present** | CinY | DurY | Not essential |
| **Not present** | CinZ | DurZ | Not essential |
| **Not present** | Cinorf8 | Durorf8 | Not essential |
| **Not present** | Cinorf9 | Not present | Not essential |
| **KyaR (216aa)** | CinR (216aa) | DurR (216aa) | Regulation |
| **KyaK (372aa)** | CinK (354aa) | DurK (349aa) | Regulation |
| **KyaL (226aa)** | CinL (236aa) | DurL (235aa) | Immunity |
| **Kyaorf11 (295aa)** | Cinorf11 (396aa) | Durorf11 (396aa) | Not essential |
| **KyaR1 (260aa)** | CinR1 (261aa) | DurR1 (261aa) | Regulation |

To detect production of kyamicin we grew all three strains on a range of 13 liquid media (Table S2) and extracted after four, five, six and seven days of growth, using (individually) methanol and ethyl acetate. Analysis of the extracts using UPLC/MS failed to identify the anticipated product (the methods were validated using authentic duramycin). This was consistent with parallel bioassays which failed to show any antibacterial activity for the extracts against *Bacillus subtilis* EC1524, which is sensitive to cinnamycin (24). Similarly, no activity was observed in overlay bioassays.

### Activation of the kyamicin BGC

Cinnamycin production and self-immunity ultimately rely on two gene products (38). The transcription of the biosynthetic genes is driven by CinR1, a SARP (*Streptomyces* Antibiotic Regulatory Protein, which usually act as pathway specific transcription activators), and self-immunity is conferred by a methyl transferase (Cinorf10) that modifies PE in the membrane to prevent binding of cinnamycin. We reasoned that transcription of the homologues of these two genes (*kyaR1* and *kyaL*, respectively), driven by a constitutive promoter, would circumvent the natural regulatory mechanism and initiate production of kyamicin. To achieve this, we made a synthetic construct, pEVK1, containing *kyaR1-kyaL* (in that order), with a *NdeI* site overlapping the start codon of *kyaR1*, a HindIII site after the stop codon of the *kyaL* and with the *kyaN* ribosome binding site (RBS) located between the two genes (the *kyaN* RBS was chosen as its sequence is most similar in the BGC to that of an ideal RBS) (Fig. S2A). The *kyaR1-kyaL* cassette was cloned into pGP9 (39) to yield pEVK4 which was introduced into the three *Saccharopolyspora* strains by conjugation. This resulted in single copies of the plasmid integrated at the *φ*BT1 phage integration site of each strain. Exconjugants were assayed by overlaying with *B. subtilis* EC1524, revealing zones of clearing for all three strains containing pEVK4 (Fig. 2A, Fig. S3). For the KY21 ex-conjugant, agar plugs were taken from the zone of clearing, extracted with 5% formic acid and analysed by UPLC/MS (Fig. 2A). In contrast to the relevant controls, an ion at *m/z* 899.36 was observed corresponding to the expected [M + 2H]^2+^ ion of kyamicin (Table 2).

**FIG 2.**
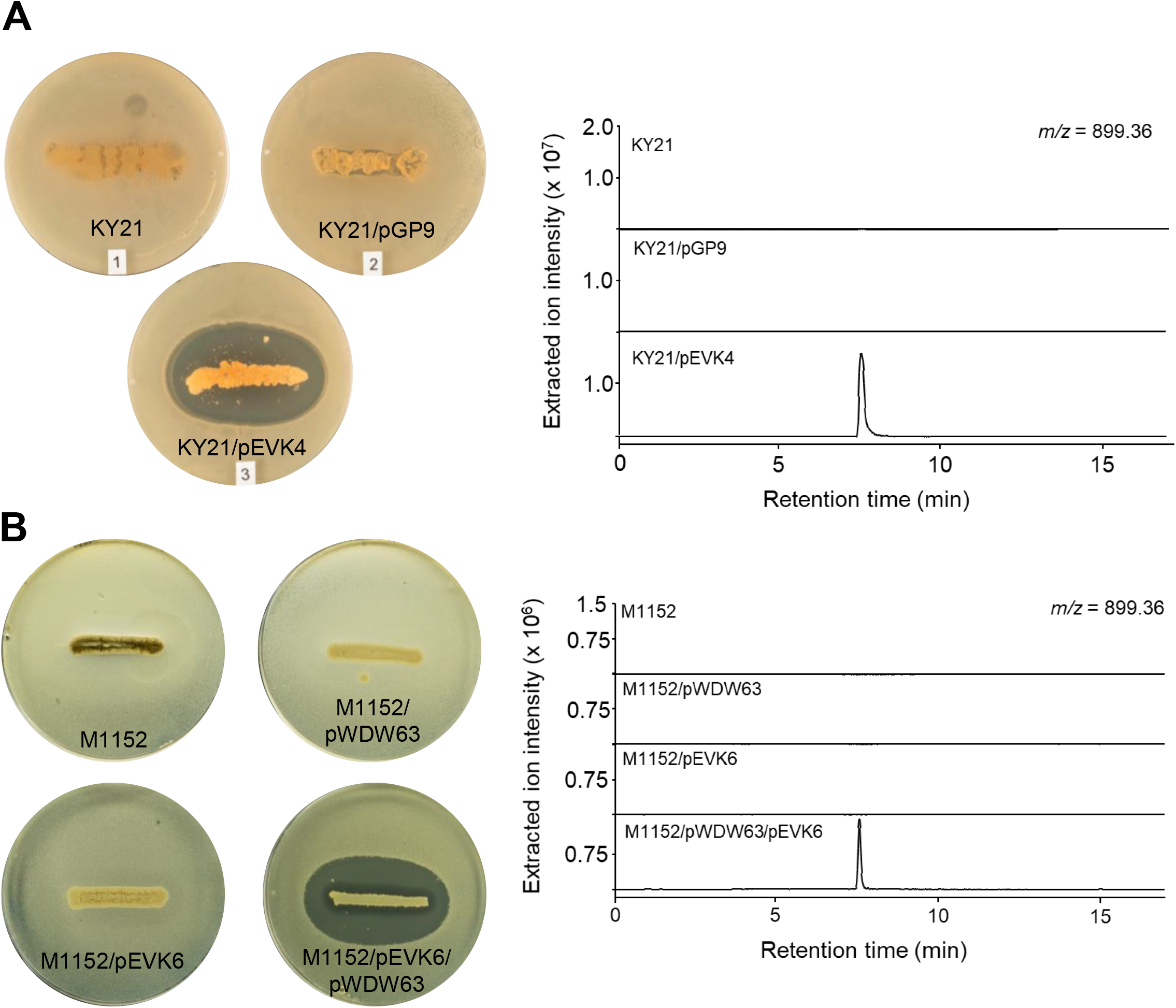
Activation of kyamicin biosynthesis and heterologous expression. Overlay bioassays were carried out with *B. subtilis* EC1524 and agar plugs were taken adjacent to the central streak and analysed by UPLC/MS. Extracted ion chromatograms are shown where *m/z* = 899.36 ([M + 2H]^2+^). Images and LC traces are representative of at least three biological repeats. **(A)** Activation of kyamicin production in KY21 strains. The pEVK4 vector containing *kyaR1* and *kyaL* results in a zone of inhibition, corresponding to the production of kyamicin, in contrast to the pGP9 empty vector control or the wildtype strain. **(B)** Heterologous expression of kyamicin in *S*. *coelicolor* M1152. A zone of inhibition, corresponding to kyamicin production, is observed only when the pWDW63 carrying the *kya* biosynthetic genes is expressed in combination with pEVK6 carrying *kyaR1* and *kyaL*.

**TABLE 2:** Calculated and observed *m/z* values for lantibiotic compounds in this study.

| Compound | Formula | Calculated<br>[M + 2H]<sup>2+</sup> $m/z$ | Observed<br>[M + 2H]<sup>2+</sup> $m/z$ | Error (ppm) |
| --- | --- | --- | --- | --- |
| Kyamicin | C<sub>76</sub>H<sub>108</sub>N<sub>20</sub>O<sub>25</sub>S<sub>3</sub> | 899.3551 | 899.3553 | 0.22 |
| Deoxykyamicin | C<sub>76</sub>H<sub>108</sub>N<sub>20</sub>O<sub>24</sub>S<sub>3</sub> | 891.3576 | 891.3557 | -2.13 |
| Partially Reduced<br>Kyamicin | C<sub>76</sub>H<sub>110</sub>N<sub>20</sub>O<sub>25</sub>S<sub>2</sub> | 884.3768 | 884.3767 | -0.11 |
| Partially Reduced<br>Kyamicin | C<sub>76</sub>H<sub>112</sub>N<sub>20</sub>O<sub>25</sub>S | 869.3987 | 869.3990 | 0.35 |
| Reduced Kyamicin | C<sub>76</sub>H<sub>114</sub>N<sub>20</sub>O<sub>25</sub> | 854.4204 | 854.4202 | -0.23 |
| Duramycin | C<sub>89</sub>H<sub>125</sub>N<sub>23</sub>O<sub>25</sub>S<sub>3</sub> | 1006.9262 | 1006.9232 | -2.98 |
| Deoxyduramycin | C<sub>89</sub>H<sub>125</sub>N<sub>23</sub>O<sub>24</sub>S<sub>3</sub> | 998.9287 | 998.9253 | -3.40 |

### Heterologous expression of the kyamicin BGC

Attempts to scale up cultures of *Saccharopolyspora* sp. KY21/pEVK4 to generate sufficient material for further study were not successful. Consequently, we attempted heterologous expression of the *kya* BGC in the well-established host *Streptomyces coelicolor* M1152 (40). To achieve this, we cloned *kyaR1L* as a *NdeI/HindIII* fragment into pIJ10257, a *φ*BT1-based integrative expression vector with a hygromycin resistance marker (41); this yielded pEVK6, which has the constitutive *ermE*\* promoter driving expression of *kyaR1L*. We then commissioned a synthetic operon containing *kyaN-H* plus the upstream promoter region of *kyaN* as an *EcoRI/XbaI* fragment (Fig. S2B). This was cloned into pSET152 (42) to give pWDW63, which integrates into the *S*. *coelicolor* chromosome at the *φ*C31 integration site, conferring apramycin resistance. pEVK6 and pWDW63 were then introduced sequentially into *S*. *coelicolor* M1152 *via* conjugation, and apramycin plus hygromycin resistant ex-conjugants were grown on R5 agar and overlaid with *B. subtilis* EC1524. In contrast to the control strains, these gave a pronounced zone of clearing. Agar plugs were taken from the zone of clearing, extracted and analysed by UPLC/MS, revealing the expected [M + 2H]^2+^ ion for kyamicin which was not present in the controls (Fig. 2B). In addition to kyamicin, a second minor new compound was observed with an *m/z* value of 891.36, consistent with the production of a small amount of deoxykyamicin presumably reflecting incomplete β-hydroxylation of the aspartic acid residue at position 15 (Table 2 and Fig. S4).

Having established the production of kyamicin in the M1152 heterologous host, we used this system to better understand how each gene product contributes to the activation of kyamicin biosynthesis. We cloned *kyaL* and *kyaR1* separately into pIJ10257, to give pEVK12 and pEVK13, respectively. Each plasmid was then introduced into M1152 alongside pWDW63, and doubly antibiotic resistant ex-conjugants were selected. These were grown on R5 agar plates and overlaid with *B*. *subtilis* EC1524; agar plugs were extracted from the resulting bioassay plates as before. For M1152/pEVK12 (*kyaL* only) no growth inhibition of the bioassay strain was observed and we could not detect kyamicin or deoxykyamicin using UPLC/MS. For M1152/pEVK13 (*kyaR1* only), we observed a zone of inhibition which was approximately three times smaller than for the M1152/pEVK6 (*kyaR1L*) positive control. UPLC/MS analysis of the M1152/pEVK13 strain detected only deoxykyamicin (Fig. S4). This is consistent with previous work which reported that deoxy versions of lantibiotics have lower biological activity (43).

### Isolation, structure elucidation and bioactivity

To isolate and verify the structure of kyamicin, growth of *S*. *coelicolor* M1152/pEVK6/pWDW63 was scaled up in liquid culture and the cell pellet extracted with 50% methanol. Crude extracts were further purified using semi-preparative HPLC to yield pure kyamicin (2.5 mg).

As the methyllanthionine bridges of kyamicin limit the ability to induce fragmentation in MS/MS experiments, the lantibiotic was subjected to chemical reduction with NaBH_4_-NiCl_2_ using a procedure published previously for the related molecule cinnamycin B (28). This leads to removal of the methyllanthionine bridges and, as anticipated, UPLC/MS of the product molecule showed an [M + 2H]^2+^ ion at *m/z* 854.42 corresponding to the loss of three sulfur atoms and gain of six hydrogen atoms (Table 2 and Fig. 3). Tandem MS experiments were carried out using both ESI and MALDI-ToF methods. Whilst ESI gave a complex mixture of fragmentation ions, for MALDI-ToF the *y* ion (NH_3_^+^) series could be clearly observed, with fragmentation at the lysinoalanine bridge appearing to occur via a rearrangement to give a glycine residue at position 6 and N=CH_2_ at the end of the lysine side chain (Fig. S5). The connectivity of the peptide was consistent with the primary sequence of kyamicin predicted by our bioinformatics analysis.

**FIG 3.**
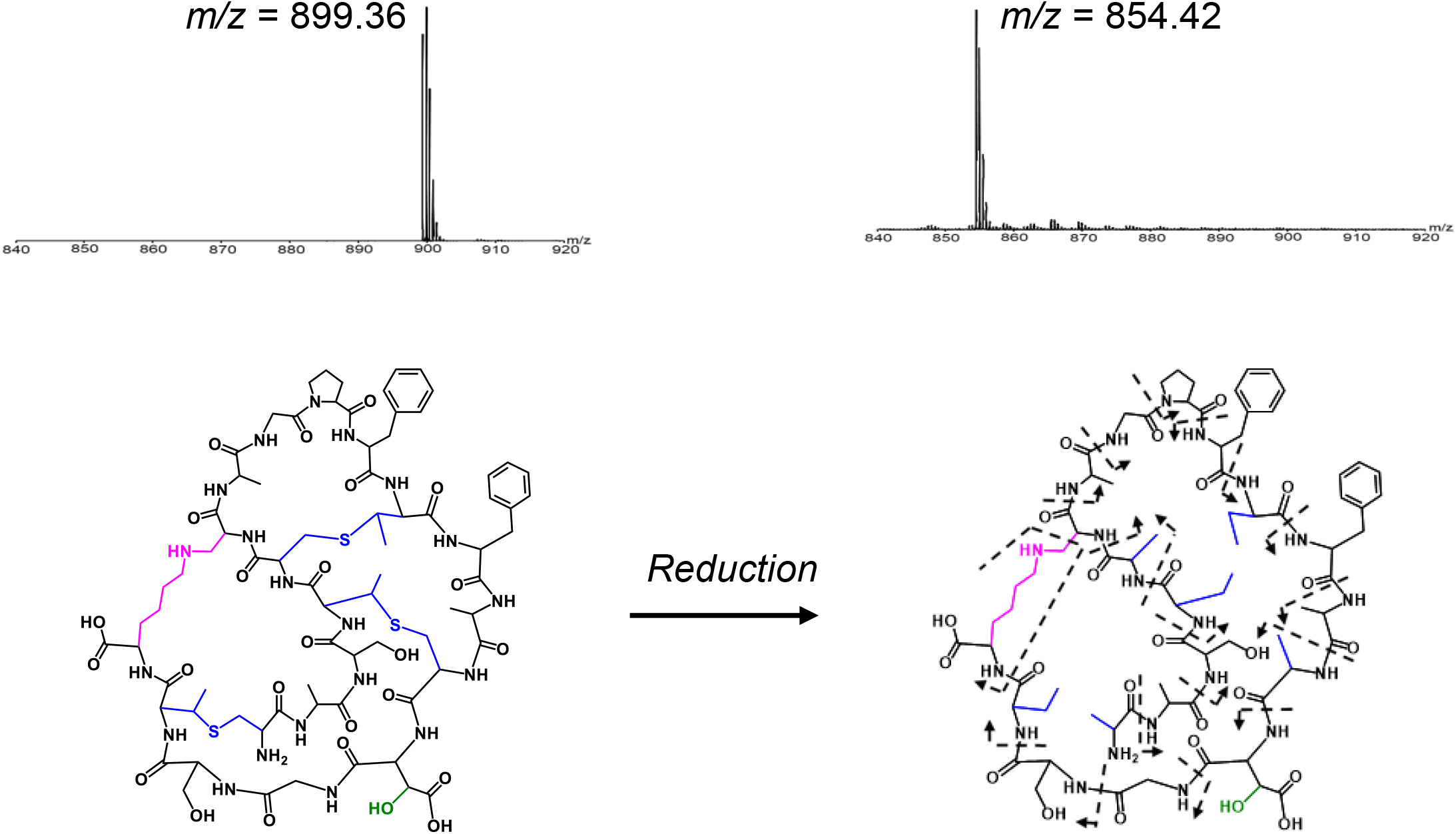
Characterisation of kyamicin. The connectivity of the peptide was confirmed by chemical reduction followed by tandem MS fragmentation. Reduction with NaBH_4_-NiCl_2_ resulted in the cleavage of the methyllanthionine bridges (blue), corresponding to the loss of three S atoms and gain of six H atoms, with a mass shift from [M + 2H]^2+^ = 899.36 *m/z* to 854.42 *m/z*. Tandem MS using the MALDI-ToF LIFT method allowed identification of the *y* ion (NH_3_^+^) series for the complete peptide (Figure S5). Fragmentation of the lysinoalanine bridge (pink) occurred via rearrangement to give N=CH_2_ at the terminus of the lysine sidechain and a glycine residue at position 6.

The chemical structure was further examined by NMR experiments comprising ^1^H, HSQC, TOCSY and NOESY analyses. Overall, 14 spin systems could be partially or completely identified in the TOCSY spectrum. These could be putatively assigned based on their spatial relationship determined from the NOESY spectrum. Coupling in the HSQC spectrum then allowed identification of several C atoms in the molecule. Spectra and assignments can be found in Fig. S6 and Table S3.

The bioactivity of the purified compound was compared with cinnamycin and duramycin using the spot-on-lawn method. The minimum inhibitory concentration (MIC) of kyamicin against *B. subtilis* EC1524 was 128 μg/mL, whereas duramycin inhibited at 32 μg/mL and cinnamycin at 16 μg/mL, representing a 4 and 8-fold MIC increase respectively (Fig. 4).

**FIG 4.**
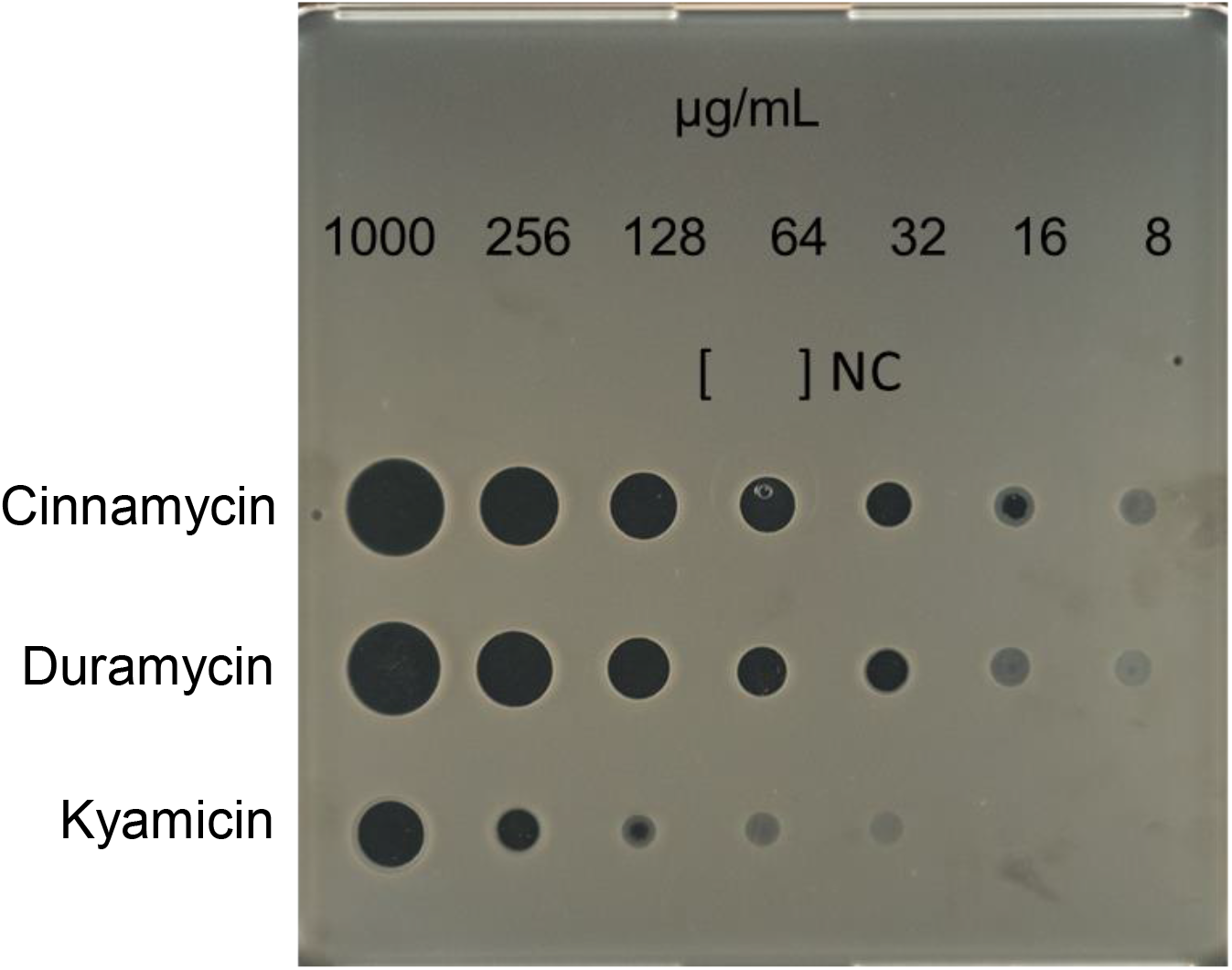
Comparative bioassay of kyamicin, duramycin and cinnamycin against *B. subtilis* EC1524. The MIC of each substance was determined by direct application of serial dilutions of the compounds in water, on a SNA agar plate inoculated with *B. subtilis* EC1524. NC = H_2_O is the negative control. Kyamicin displays an MIC of 128 μg/mL, whereas duramycin inhibits at 32 μg/mL and cinnamycin at 16 μg/mL.

### Cross species activation of the duramycin BGC

Many cinnamycin-like BGCs can be identified in the published sequence databases, but their products remain cryptic. Thus, the potential of the *kyaR1-kyaL* construct to induce expression of other cinnamycin-like lantibiotics was explored.

The BGC for duramycin was cloned previously from *Streptomyces cinnamoneus* ATCC 12686 (Fig. S7A) but attempts to produce the lantibiotic heterologously failed. Consequently, the duramycin BGC was reconfigured in pOJKKH, which contains all the biosynthetic genes, but lacks immunity and regulatory genes, and has a SARP binding site upstream of *durN* that is similar to that upstream of *kyaN* (Fig. S7B) (38). pOJKKH and pEVK6 were introduced sequentially into *S*. *coelicolor* M1152 *via* conjugation and the resulting ex-conjugants assessed for duramycin production. Overlay bioassays using *B. subtilis* EC1524 indicated the production of an antibacterial molecule by *S*. *coelicolor* M1152/pOJKKH/pEVK6 (Fig. 5). Agar within the growth inhibition zone was extracted and the resulting sample analysed by UPLC/MS. An ion at *m/z* 1006.92 was observed, corresponding to the expected [M + 2H]^2+^ ion for duramycin (Table 2). The production of duramycin was confirmed by comparison to an authentic standard. A deoxy derivative was also detected with an *m/z* of 998.93 (Table 2), typically at ~30% the level of duramycin. Expression of pOJKKH alone or in conjunction with the empty pIJ10257 vector did not result in duramycin biosynthesis, demonstrating that expression of both *kyaR1* and *kyaL* are required to induce heterologous duramycin biosynthesis in *S*. *coelicolor* M1152. Thus, we have shown that the SARP and resistance genes from a cinnamycin-like BGC from a *Saccharopolyspora* species can be used to activate a cinnamycin-like BGC from a *Streptomyces* species, a cross genus activation.

**FIG 5.**
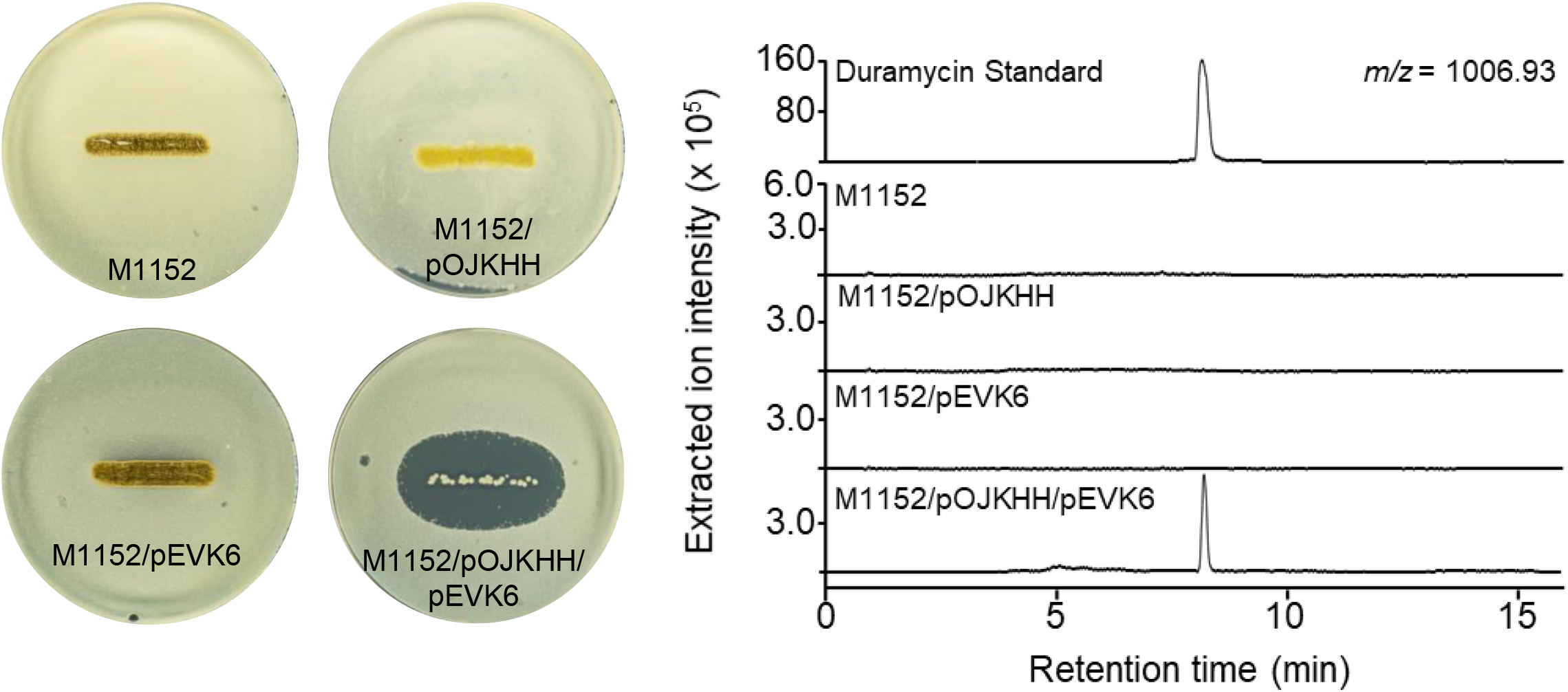
Activation of duramycin biosynthesis. Overlay bioassays were carried out with *B. subtilis* EC1524 and agar plugs were taken adjacent to the central streak and analysed by UPLC/MS. Extracted ion chromatograms are shown where *m/z* = 1006.93 ([M+2H]^2+^). Duramycin was only detected in the strain carrying both pOJKHH and pEVK6. The duramycin peak aligns with an authentic standard of duramycin (1 mg/mL in 5% formic acid), shown on a separate scale. Images and LC traces are representative of at least three biological repeats.

## DISCUSSION

Three isolates from the relatively rare actinomycete genus *Saccharopolyspora* were isolated from the external microbiome of *T. penzigi* plant ants collected at two locations in Kenya more than 50 km apart (13). Despite this geographical separation, their genomes were extremely similar and analysis using antiSMASH identified almost identical biosynthetic capabilities. Amongst the conserved BGCs was one encoding a cinnamycin-like lantibiotic which we named kyamicin.

Despite culturing on a wide range of media, we were unable to elicit production of kyamicin in the wild-type *Saccharopolyspora* strains. The production of cinnamycin in *S*. *cinnamoneus* DSM 40005 requires the expression of two key genes, *cinR1* and *cinorf10*, encoding a pathway specific regulatory gene (a SARP) and a self-immunity gene (a PE methyltransferase), respectively (38). As the *kya* BGC encodes homologues of these genes, we expressed them constitutively in the three *Saccharopolyspora* strains which led to activation of the BGC and production of kyamicin. Since we were unable to isolate enough kyamicin from these strains for further study, a heterologous production platform was developed using *S. coelicolor* M1152 which allowed us to confirm the structure of kyamicin and assess its antibacterial activity.

Having demonstrated the utility of a constitutively expressed SARP/self-immunity cassette for driving expression of the otherwise silent *kya* BGC we utilised this knowledge to activate duramycin production in a heterologous host. Contemporaneous with our experiments, the duramycin BGC was also identified by genome sequencing of *S*. *cinnamoneus* ATCC 12686 (33). This analysis described the same genomic region containing *durN* to *durH* and surrounding genes (Table 1) but failed to reveal putative regulatory and immunity genes. Co-expression of *durA, durM*, *durN* and *durX* in *E. coli* was sufficient to direct the biosynthesis of duramycin A, and the functions of DurA, DurM, DurN and DurX were confirmed by detailed biochemical analyses. Our subsequent bioinformatic analysis of the published genome sequence identified homologs of the resistance genes *cinorf10/kyaL* and the regulatory genes *cinRKR1/kyaRKR1* in region 54637 to 59121 bp of contig MOEP01000113.1 from the deposited genome sequence (accession no. NZ_MOEP00000000). This region is separated from the *dur* biosynthetic genes by a section of low mol %GC DNA, the analysis of which suggests that a phage or other mobile element may have inserted between *durZ* and *durorf8* (Fig. S7). Thus, it appears likely that the immunity and regulatory mechanisms described previously for the control of cinnamycin biosynthesis are conserved for duramycin biosynthesis in *S*. *cinnamoneus* ATCC 12686.

Given the potential utility of cinnamycin-like class II lanthipeptides in several therapeutic contexts, the ability to generate analogues of these compounds with modified properties and in sufficient quantity for preclinical assessment is of significant value. The methods described here provide a platform for the identification of additional natural lanthipeptides whose biosynthesis cannot be detected in the host strain, and for the diversification of their chemical structures to generate new-to-nature molecules.

## MATERIALS AND METHODS

### Bacterial strains, plasmids and growth conditions

All bacterial strains and plasmids used in this study are listed in Table S1 in the supplemental material. *Saccharopolyspora* and *Streptomyces* strains were grown on soya flour mannitol (SFM) agar medium with appropriate antibiotics at 30 °C unless otherwise stated. *E. coli* and *B. subtilis* EC1524 strains were grown on lysogeny broth (LB) medium with appropriate antibiotics at 37 °C. R5 agar (44) was used for bioassay plates.

### DNA extraction and genomic analysis

The salting out method (44) was used to extract genomic DNA. The DNA was sequenced at the Earlham Institute (Norwich, UK) using SMRT sequencing technology (Pacific Biosciences RSII platform) and assembled using the HGAP2 pipeline (36).

### Overlay bioassays

For each strain to be tested, a streak from a spore stock was applied in the centre of an R5 agar plate and left to grow for seven days. *B. subtilis* EC1524 was grown from a single colony overnight, then diluted 1:20 in fresh media and grown until OD_600_ = 0.4 − 0.6. The exponential culture was mixed with 1:100 molten soft nutrient agar (SNA) (44) and the mixture was used to overlay the plate (5 mL SNA mixture/agar plate). The plate was incubated at room temperature overnight.

### Extractions from overlay bioassays

Plugs of agar 6.35 mm in diameter were taken adjacent to the streaked actinomycete strain on an overlay bioassay plate, corresponding to the zone of growth inhibition where one was observed. Agar plugs were frozen at-80 °C for 10 min, thawed and then 300 μL of 5% formic acid was added. This was vortexed briefly and shaken for 20 min. After centrifugation (15,682 × *g* for 15 min) the supernatant was collected and filtered using a filter vial (HSTL Labs) prior to UPLC-MS analysis.

### UPLC-HRMS

Data were acquired with an Acquity UPLC system (Waters) equipped with an Acquity UPLC^®^ BEH C18 column, 1.7 μm, 1×100 mm (Waters) connected to a Synapt G2-Si high-resolution mass spectrometer (Waters). For analytical UPLC 5.0 μL of each sample was injected and eluted with mobile phases A (water/0.1% formic acid) and B (acetonitrile/0.1% formic acid) at a flow rate of 80 μL/min. Initial conditions were 1% B for 1.0 min, ramped to 40 % B within 9.0 min, ramped to 99 % B within 1.0 min, held for 2 min, returned to 1 % B within 0.1 min and held for 4.9 min.

MS spectra were acquired with a scan time of 1.0 s in the range of *m/z* = 50 − 2000 in positive resolution mode. The following parameters were used: capillary voltage of 3.0 kV, cone voltage 40 V, source offset 80 V, source temperature 130 °C, desolvation temperature 350 °C, desolvation gas flow of 700 L/h. A solution of sodium formate was used for calibration. Leucine encephalin peptide (H_2_O/MeOH/formic acid: 49.95/49.95/0.1) was used as lock mass (556.2766 *m/z*) and was injected every 30 s during each run. The lock mass correction was applied during data analysis.

### Design of *kya* BGC activation and immunity plasmids

pEVK1, a pUC57 derivative, contains the synthetic *kyaR1* and *kyaL* (Genscript) arranged as an operon. pEKV1 has a *NdeI* site overlapping the start codon of *kyaR1* and a HindIII site immediately after the stop codon of *kyaL* with the two genes separated by a short intergenic region containing a RBS designed from the RBS of *cinN* (Fig. S2A). The *NdeI-HindIII kyaR1L* fragment from pEVK1 was cloned in pGP9 (39) to give pEVK4, and into pIJ10257 (41) to give pEKV6. *kyaR1* and *kyaL* were amplified individually as *NdeI-HindIII* compatible fragments using the primers AmplkyaR1-F (GCGCAAGCTTCTACGACGCGGTGTGA) and AmplkyaR1-R (GCGCGCCATATGAAACCGCTGTCGTTCC) for *kyaR1*, and AmplkyaL-F (GCGCGCCAT AT GGAT CCAGT ACAGACCA) and AmplkyaL-R (GCGCAAGCTTTCAGCGGTCCTCCGCC) for *kyaL;* they were cloned as *NdeI-HindIII* fragments into pIJ10257 to yield pEVK12 and pEVK13 respectively. PCR generated fragments were verified by Sanger sequencing.

### Cloning the duramycin BGC from *Streptomyces cinnamoneus* ATCC 12686

The cloning of a ~5 Kb *Bg/II* fragment of chromosomal DNA to create pIJ 10100 was described previously (24). This plasmid has a *KpnI* site in the middle of *durX*. *KpnI* fragments upstream and downstream of this *KpnI* site were identified by Southern blotting and isolated by creating a mini-library of *KpnI* fragments in pBluescriptIIKS followed by colony hybridization to give pDWCC2 and pDWCC3, respectively. Analysis of the sequence of these plasmids identified 15 genes (shown in Fig. S7). A plasmid carrying the duramycin biosynthetic genes but not the putative phage DNA was prepared by digesting pDWCC3 with *XhoI* and HindIII (site is in the multiple cloning site of pBluescriptIIKS) removing the 5’ end of *durZ* and the putative phage DNA. This region was replaced with a *XhoI* and HindIII cut PCR fragment that reconstituted the portion of *durZ* removed in the previous step and introduced a *HindIII* site upstream of the *durZ* start codon. The 666 bp PCR fragment was generated using the primers BK10 (GAGCTTGACGCCGCCGAAGTAGC) and Hindprim (GCGGCGAAGCTTGAGGTGGCCTCCTCCACGAAGCCA) with pDWCC3 as template and was cut with *XhoI* plus *HindIII* to give a 363 bp fragment. The resulting plasmid was then digested with *KpnI* plus *XbaI* (the *XbaI* site is in the multiple cloning site of pBluescriptIIKS) and the fragment carrying putative duramycin genes was cloned into *KpnI* plus *XbaI* cleaved pOJ436 to give pOJKH. The *KpnI* fragment from pDWCC2 was then cloned into pOJKH cut with *KpnI* to give pOJKKH which was verified by *Bg/II* digestion, thus restoring the original gene context.

### Isolation and purification of kyamicin

*S*. *coelicolor* M1152/pWDW63/pEVK6 was grown in tryptic soy broth (12 × 500 mL in 2.5 L Erlenmeyer flasks) and incubated at 28 °C and 200 rpm on an orbital shaker for seven days. The cells were harvested and extracted with methanol/water (1:1; 500 mL) with ultrasonication for 2 h and subsequent shaking for 16 h. After centrifugation, the supernatant was filtered and concentrated under vacuum giving 613 mg of crude material, which was then purified by semi-preparative HPLC. Chromatography was achieved over a Phenomenex Gemini-NX reversed-phase column (C18, 110 Å, 150 × 21.2 mm) using a Thermo Scientific Dionex UltiMate 3000 HPLC system. A gradient was used with mobile phases of A: H_2_O (0.1% formic acid) and B: methanol; 0–1 min 10% B, 1–35 min 10-85% B, 35–40 min 85–100% B, 40–45 min 100% B, 45–45.1 min 100–10% B, 45.1-50 min 10% B; flowrate 20 mL/min; injection volume 1000 μL. Absorbance was monitored at 215 nm and fractions (20 mL) were collected and analysed by UPLC/MS. Kyamicin was observed in fractions 22-25 which were combined and concentrated to yield an off-white solid (2.5 mg).

### Minimum inhibitory concentration (MIC) determination

The spot-on-lawn method was used to determine lantibiotic MICs. A 1000 μg/mL stock solution of each lantibiotic was prepared using sterile water, along with serial dilutions from 256 – 8 μg/mL. *B. subtilis* EC1524 was grown and mixed with molten SNA as described above to create a lawn of bacterial growth. Once set, 5 μL of each dilution was applied directly to the agar and incubated overnight at room temperature. The MIC was defined as the lowest concentration for which a clear zone of inhibition was observed.

### Chemical reduction of kyamicin

Kyamicin (1 mg) was dissolved in methanol (0.5 mL) and added to an aqueous solution of NiCl_2_ (20 mg/mL; 0.5 mL). The solution was mixed with solid NaBH_4_ (5 mg), resulting in the generation of hydrogen gas and the formation of a black Ni_2_B precipitate. The tube was immediately sealed, and the mixture stirred at 55 °C. The reaction progress was monitored by UPLC-HRMS as described above, for which a peak with an *m/z* of 899.36 was observed for kyamicin ([M + 2H]^2+^). The successive formation of peaks with the following masses were observed: *m/z* = 884.38, 869.40 and 854.42, corresponding to the successive reduction of the three thioether bridges. After 5 h only the ion with *m/z* 854.42 could be observed, indicating that the starting material had been completely reduced. The precipitate was collected by centrifugation at 15,682 × *g* for 10 min. As the reaction supernatant contained only trace amounts of the desired product, a fresh solution of MeOH/H_2_O 1:1 (0.5 mL) was added to the precipitate and it was subject to ultrasonication for 30 min. Reduced kyamicin was then detected in sufficient quantity for MS/MS experiments to confirm the peptide sequence.

### MS analysis of reduced kyamicin

For ESI/MS^2^ analysis the mass of interest (854.42) was selected using an inclusion list and fragmented using data directed analysis (DDA) with the following parameters: top3 precursor selection (inclusion list only); MS2 threshold: 50,000; scan time 0.5 s without dynamic exclusion. Collision energy (CE) was ramped between 15-20 at low mass (50 *m/z*) and 40-100 at high mass (2000 *m/z*). Further increase of the CE to 20-30/60-120 led to complete fragmentation.

For MALDI-ToF/MS the samples were mixed with α-cyano-4-hydroxycinnamic acid as matrix and analysed on an AutoflexTM Speed MALDI-TOF/TOF mass spectrometer (Bruker DaltonicsTM GmbH). The instrument was controlled by a flexControlTM (version 3.4, Bruker) method optimised for peptide detection and calibrated using peptide standards (Bruker). For sequence analysis fragments produced by PSD were measured using the LIFT method (Bruker). All spectra were processed in flexAnalysisTM (version 3.4, Bruker).

### NMR experiments

NMR measurements were performed on a Bruker Avance III 800 MHz spectrometer. Chemical shifts are reported in parts per million (ppm) relative to the solvent residual peak of DMSO-d_6_ (^1^H: 2.50 ppm, quintet; ^13^C: 39.52 ppm, septet).

## ACKNOWLEDGEMENTS

This work was supported by the Biotechnology and Biological Sciences Research Council (BBSRC) *via* Institute Strategic Program BB/P012523/1 to the John Innes Centre (JIC); by Research Grant BB/P021506/1 to B.W.; by Research Grant 208/P08242 to M.J.B; and by NPRONET Proof of Concept Award BB/L013754/1 to B.W and M.I.H. It was also supported by Research Grants NE/M015033/1 and NE/M014657/1 from the Natural Environment Research Council (NERC) to M.I.H. and B.W. E.V.K. was supported by a Norwich Research Park (NRP) studentship and the BBSRC NRP Doctoral Training Partnership grant BB/M011216/1. We acknowledge the Earlham Institute (Norwich, UK) for sequencing and assembly of the *Saccharopolyspora* sp. KY3, KY7 and KY21 genomes, which was funded by a Norwich Research Park Translational Award to B.W. and M.I.H. D.J.M.’s work on ants in Kenya is supported by the National Geographic Society, Nature Kenya and the National Commission of Science Technology and Innovation (NACOSTI Permit # MOST13/001/35C136). Dr. Juan Pablo Gomez-Escribano (JIC) is thanked for his valuable input with analysis of the *Saccharopolyspora* sp. KY3, KY7 and KY21 genomes. We thank Dr Lionel Hill and Dr Gerhard Saalbach (JIC) for excellent metabolomics support. Dr. Jesus Angulo and Dr Ridvan Nepravishta (UEA) are thanked for their assistance with NMR data acquisition. The authors declare no competing financial interests.

## SUPPLEMENTAL MATERIAL

**FIG S1 Alignment of *Saccharopolyspora* sp. KY3, KY7 and KY21 16S rDNA sequences**. The alignment was performed with Clustal Omega (v1.2.4) and the figure was generated by SnapGene Viewer (v4.2.11). The difference between KY21 to strains KY3 and KY7 is indicated with a black arrow and a box at position 685.

**FIG S2 Schematic of synthetic artificial operons**. **(A)** The operon consisting of *kyaR1*, encoding a *Streptomyces* antibiotic regulatory protein (SARP), and *kyaL*, encoding a PE-methyl transferase that provides resistance – the homologues of *cinR1* and *cinorf10* respectively. **(B)** The operon carrying genes *kyaN* to *kyaH* as an *EcoRI/Xbal* fragment. These genes are expected to be essential for kyamicin biosynthesis.

**FIG S3 Activation of kyamicin biosynthesis in KY3 and KY7**. The pEVK4 vector containing *kyaR1* and *kyaL* results in a zone of inhibition, corresponding to the production of kyamicin, in contrast to the pGP9 empty vector control or the wildtype strain. **(A)** Activation of kyamicin production in KY3, and **(B)** in KY7.

**FIG S4 Dissection of the contribution of *kyaR1* and *kyaL* to kyamicin BGC activation**. Overlay bioassays were carried out with *B. subtilis* EC1524 and agar plugs were taken adjacent to the central streak and analysed by UPLC/MS. Expression of *kyaL* (pEVK12) does not result in a zone of inhibition. Expression of *kyaR1* (pEVK1) results in a zone of inhibition, corresponding to the production of deoxykyamicin only. Co-expression of *kyaR1* and *kyaL* (pEVK6) results in a zone of inhibition, corresponding to the production of both kyamicin and deoxykyamicin. Images and LC traces are representative of at least three biological repeats. **(A)** Extracted ion chromatograms are shown where *m/z* = 899.36 ([M+2H]^2+^). **(B)** Extracted ion chromatograms are shown where *m/z* = 891.36 ([M+2H]^2+^).

**FIG S5 Kyamicin fragmentation**. Following reduction to remove methyllanthionine bridges, kyamicin was subject to MALDI-ToF tandem MS, giving the complete *y* ion (NH_3_^+^) series. **(A)** Structure of reduced kyamicin and the *y*_1_ – *y*_18_ ion series. **(B)** MALDI-ToF tandem MS spectrum with the *y* ion series indicated with dashed red lines.

**FIG S6 Kyamicin NMR Spectra**. **(A)** ^1^H NMR spectrum. **(B)** TOCSY spectrum. **(C)** NOESY spectrum. **(D)** HSQC spectrum.

**FIG S7 Schematic of duramycin BGC and plasmids used to construct pOJKKH and SARP binding sites of kyamicin, cinnamycin and duramycin**. **(A)** The *S*. *cinnamoneus* DNA sequences represented on the plasmids pDWCC2 and pDWCC3 are present in the published genome sequence as 81593-99144 bp of contig NZ_MOEP01000024.1. pDWCC2 consists of the area from the left side *KpnI* site (from *durorfl*) to the central side *KpnI* site in *durX*. pDWCC3 consists of the area covering from the central *KpnI* site in *durX* to the right side *KpnI* site after a putative integrase encoding gene. The putative duramycin resistance/regulatory genes are represented in the published genome sequence by 54637-59121 bp of contig NZ_MOEP01000113.1. **(B)** Sequence alignment of putative SARP binding sites of kyamicin, cinnamycin and duramycin. Conserved residues within all three sequences are marked with asterisks and the 5 bp SARP binding motifs are in bold. The alignment was performed with Clustal Omega (v1.2.4).

**TABLE S1 Strains and plasmids used in this work**.

**TABLE S2 Recipes for liquid screening media**. Quantities of components are given in g/L. SM = screening media.

**TABLE S3 Putative NMR assignments**. ND = not determined.

